# Vibrotactile auricular vagus nerve stimulation alters limbic system connectivity in humans: A pilot study

**DOI:** 10.1101/2024.09.10.612366

**Authors:** Kara M. Donovan, Joshua D. Adams, Ki Yun Park, Phillip Demarest, Gansheng Tan, Jon T. Willie, Peter Brunner, Jenna L. Gorlewicz, Eric C. Leuthardt

**Author notes:** Corresponding authors: (KMD), (ECL). These authors contributed equally to this work and share senior authorship (Peter Brunner, Jenna L. Gorlewicz, Eric C. Leuthardt).

## Abstract

Vibration offers a potential alternative modality for transcutaneous auricular vagus nerve stimulation (taVNS). However, mechanisms of action are not well-defined. The goal of this study was to evaluate the potential of vibrotactile stimulation as a method for activating central brain regions akin to other vagal nerve stimulation methodologies. To do so, intracranial electrophysiological signals were recorded in human subjects to perform a parametric characterization of vibrotactile taVNS and investigate changes in coherence across key brain regions. We hypothesized that vibrotactile taVNS would increase coherence between limbic brain areas, similar to areas activated by classic electrical VNS approaches. Our specific regions of interest included the orbitofrontal cortex, anterior cingulate cortex, amygdala, hippocampus, and parahippocampal gyrus. Patients with intractable epilepsy undergoing stereotactic electroencephalography (sEEG) monitoring participated in the study. Vibrotactile taVNS was administered across five vibration frequencies following a randomized stimulation on/off pattern, and sEEG signals were recorded throughout. Spectral coherence in response to stimulation was defined across four canonical frequency bands, theta, alpha, beta, and broadband gamma. At the group level, vibrotactile taVNS resulted in significantly increased global low-frequency coherence. Anatomically, multiple limbic brain regions exhibited notably increased coherence during taVNS compared to baseline. The percentage of total electrode pairs demonstrating increased coherence was also quantified at the individual level. 20 Hz vibration resulted in the highest percentage of responder pairs across low-frequency coherence measures, but notable inter-subject variability was present. Overall, vibrotactile taVNS induced significant low-frequency coherence increases involving several limbic system structures. Further, parametric characterization revealed the presence of inter-subject variability in terms of identifying the optimal vibration frequency. These findings encourage continued research into vibrotactile stimulation as an alternative modality for noninvasive vagus nerve stimulation.

## 1 Introduction

Vagus nerve stimulation (VNS) has been widely investigated in both animals and humans for a multitude of applications [1–11]. To date, surgically implantable VNS has been FDA-approved for intractable epilepsy, treatment-resistant depression, and, most recently, chronic stroke [12–16]. While VNS has many beneficial effects, its invasive nature inherently limits its applications. Consequently, a noninvasive alternative, transcutaneous auricular VNS (taVNS), is also under investigation for many similar and unique indications [17–22]. The vagus nerve is a mixed fiber nerve and has afferent fibers that innervate the cymba concha and tragus regions of the outer ear, referred to as the auricular branch of the vagus nerve (ABVN) [23–26]. Importantly, current literature suggests that taVNS activates several of the same regions that are putatively believed to underly the mechanism(s) of action for implantable cervical VNS, namely, the nucleus tractus solitarius (NTS) and locus coeruleus (LC) [24,27–30]. As the primary noradrenergic nucleus in the brain, the LC produces the majority of norepinephrine (NE), which is secreted throughout the central nervous system [24,27,31]. The NTS has disynaptic projections to the LC, through which it has direct and indirect projections to several cortical and subcortical structures, such as the amygdala, hippocampus, and insula, along with other limbic system brain regions [7,24,27].

Given these substantial limbic-relevant connections, invasive and noninvasive VNS have been studied regarding their impact on various cognitive processes, such as memory and attention. Prior imaging literature has demonstrated region-specific responses to VNS in limbic areas with well-documented roles in cognition and memory function, such as the amygdala and cingulate [32–38]. Beyond just activity changes in these areas, VNS has also been shown to enhance synchronization in animal models, specifically within theta band rhythms (4-10 Hz), between the basolateral amygdala (BLA) and anterior cingulate cortex (ACC) [39]. Additionally, VNS in rodent models has been shown to increase functional connectivity between the hippocampus and retrosplenial cortex [40]. Current literature suggests that phase synchronization, also referred to as magnitude-squared coherence (MSC), across brain regions plays a role in various cognitive tasks [41–44]. Specifically, noninvasive recordings from human subjects support the notion that there are broad changes in coherence across large cortical regions during memory tasks [45]. Together, these studies suggest that VNS may modulate physiologic mechanisms and cortical and subcortical sites that are relevant to memory and other cognitive functions.

While electrical stimulation is the most common method of noninvasive taVNS, nonelectrical approaches may also activate the ABVN. Addorisio et al. demonstrated that vibrotactile stimulation administered to the cymba concha region of the outer ear could elicit vagal-mediated anti-inflammatory effects. Specifically, vibrotactile stimulation reduced inflammatory markers in healthy adults and lessened inflammatory responses in rheumatoid arthritis (RA) patients [46]. Additionally, recent work by Tan et al. revealed both arousal and working memory enhancement effects during 6 Hz vibrotactile taVNS [47]. While these studies are exciting proofs of concept, what is currently lacking is a more refined understanding of the central effects of vibrotactile taVNS in humans. Here, we performed a parametric characterization of vibrotactile taVNS, investigating direct cortical and subcortical neurophysiological responses induced by varying vibrotactile taVNS frequencies within invasively monitored human subjects. Specifically, we quantified coherence changes in several regions of interest (ROIs) – orbitofrontal cortex (OFC), ACC, amygdala, hippocampus, and parahippocampal gyrus (PHG) – using stereotactic electroencephalography (sEEG) recordings from epilepsy patients. sEEG recordings provide a unique opportunity to measure and delineate brain regions that respond to taVNS directly. ROIs were selected based on brain areas identified from rodent studies and human imaging studies, as well as brain areas that play a role in memory and cognition [6,7,24,28,29,48,49]. Based on literature indicating that VNS and taVNS have the potential to improve cognitive function, we hypothesized that vibrotactile taVNS would activate a similar pathway and thus increase neuronal communication in brain areas involved in memory [17,50,51]. In this study, we used a comprehensive coherence analysis to test our hypothesis, and our results demonstrate that specific frequencies of vibrotactile taVNS induce significant changes in limbic and memory-relevant regions similarly affected by traditional VNS approaches. These findings support the potential of vibrotactile taVNS as a useful noninvasive modality for neuromodulation.

## 2 Materials and methods

### 2.1 Participants

Nine subjects (4 males, 5 females, mean 39 ± 13 years) undergoing clinically-indicated invasive monitoring via sEEG for intractable epilepsy at Barnes Jewish Hospital participated in this study. Subjects were excluded on the basis of insufficient baseline data or large anatomical defects, resulting in seven subjects for data analysis. Within that population, 16 ± 2 electrode shanks were implanted (200 ± 29 contacts). After adjusting for recording limitations and excluding noisy contacts as well as those classified as white matter, ventricles, or unknown, an average of 174 ± 28 contacts generated good recordings for analysis. Values are reported as mean ± standard deviation unless otherwise stated. Aggregated anatomical coverage spanned 28 brain regions, with pre-determined ROIs (OFC, ACC, amygdala, hippocampus, and PHG) exhibiting dense coverage at the population level (**Fig 1**). The recruitment period for this study took place between May 16, 2022 and April 25, 2023. The study was approved by the Washington University Institutional Review Board prior to enrollment. All subjects were over 18 years of age and provided written informed consent before participating in any research procedures.

**Fig 1.**
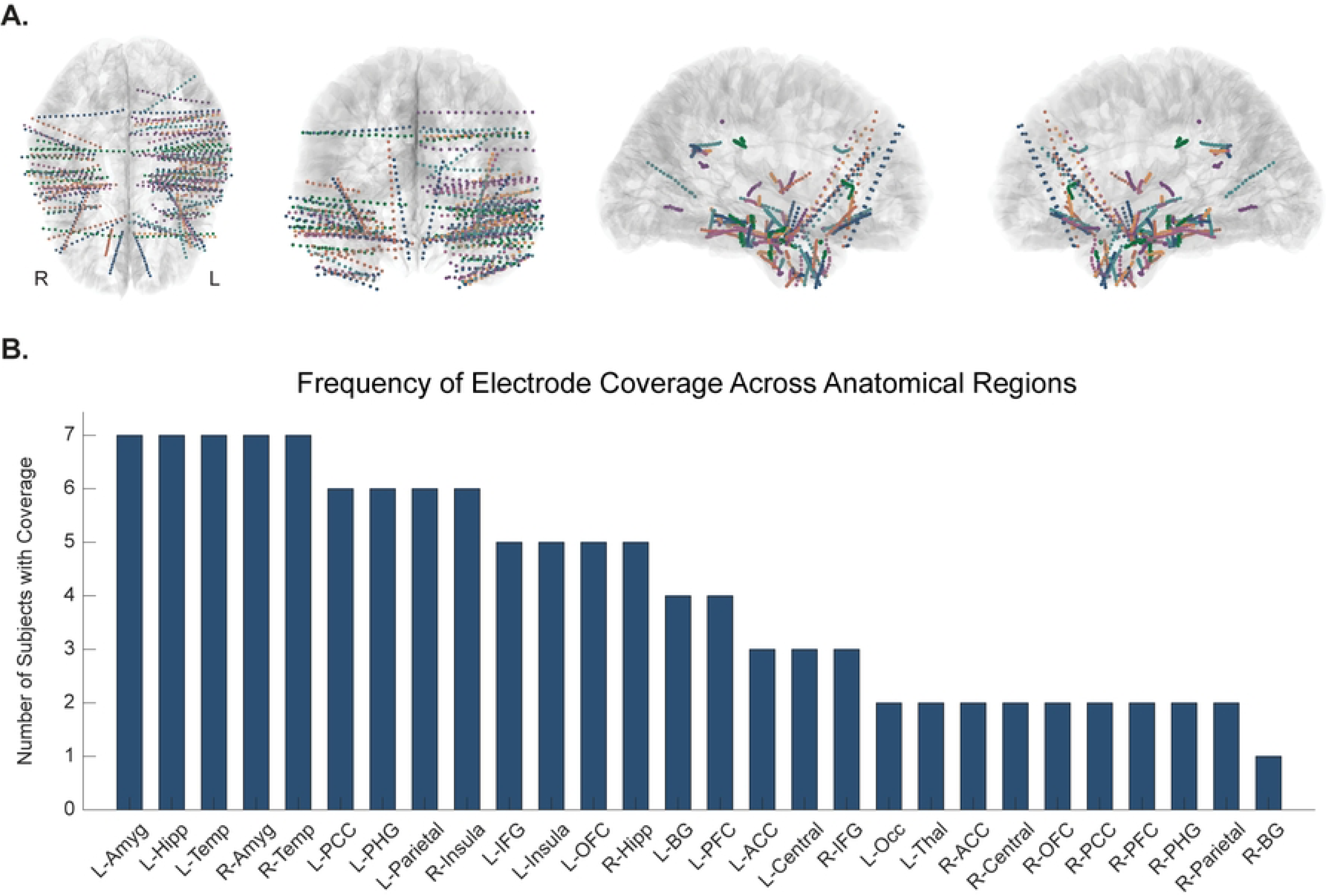
Aggregated anatomical coverage across all subjects. **(A)** Coverage of stereotactic electroencephalography (sEEG) contacts broken down by individual subjects. Each subject’s sEEG coverage is shown in a different color. Images are shown in radiological view. **(B)** sEEG contacts were classified into 28 different anatomical regions to facilitate group-level analysis. The number of subjects who had coverage in each of these anatomical regions is illustrated in the bar graph (i.e., all 7 subjects had coverage in the left amygdala, left hippocampus, left temporal lobe, right amygdala, and right temporal lobe, while only 1 subject had coverage in the right basal ganglia). ACC = anterior cingulate cortex; Amyg = amygdala; BG = basal ganglia; Hipp = hippocampus; IFG = inferior frontal gyrus; Occ = occipital lobe; OFC = orbitofrontal cortex; PCC = posterior cingulate cortex; PFC = prefrontal cortex; PHG = parahippocampal gyrus; Temp = temporal lobe; Thal = thalamus.

### 2.2 Study design

Data collection took place 1-7 days post-implantation and consisted of one experimental session lasting approximately 30 minutes. The session started with three minutes of baseline recorded at rest, followed by five-second stimulation trials alternating with five-second off periods (**S1 Fig**). Subjects remained at rest in either their hospital bed or a chair throughout the experimental session. All subjects completed 30 trials for each of five specified vibration frequencies while intracranial activity was recorded continuously.

### 2.3 Vibrotactile taVNS

#### Custom earpiece design

Because vibration is a new modality for administering taVNS, a custom vibration-delivery earpiece was designed consisting of a flexible hook backbone that wrapped around the top of the ear (**S2 Fig**) [52]. One end (the stimulating end) terminated in the cymba concha, while the other end (the stabilizing end) terminated behind the earlobe. The stimulating end consisted of a friction fit slot through which a 5 mm eccentric rotating mass (ERM) vibration motor was inserted [53]. This design was created in SolidWorks and additively manufactured in Flexible 80A material on a Formlabs Form 3B stereolithographic printer [54]. All printed components were washed in isopropyl alcohol and underwent heated post-cure per Formlabs specification prior to use. Vibration characteristics were also validated via Laser Doppler Vibrometry (LDV) benchtop testing to ensure specified frequencies were being generated.

#### Administration of taVNS

Vibrotactile taVNS was administered across five pulsed frequencies – 2, 6, 12, 20, and 40 Hz – selected to span physiologically relevant frequency bands. Based on literature suggesting neuronal responses to VNS and taVNS are temporally precise, a five-second trial duration was used [55–59]. Stimulation alternated between five seconds on, five seconds off, with each vibrotactile condition delivered for 30 trials. Trials were randomly interleaved for a total of 150 vibrotactile trials across all frequencies.

### 2.4 Data acquisition

sEEG signals were recorded at a 2,000 Hz sampling rate using the general-purpose BCI2000 software package [60]. A Nihon Kohden JE-120 A recording system (Nihon Kohden, Tokyo, Japan) was used to amplify and digitize the sEEG signals.

### 2.5 Data analysis

#### Electrode localization

Localization of sEEG electrodes was performed using the Versatile Electrode Localization Framework (VERA) [61]. The postoperative computed tomography (CT) scan was first registered to the preoperative magnetic resonance imaging (MRI) scan to generate a subject-specific brain model. Electrodes were then mapped to the FreeSurfer Desikan-Killiany atlas. To project the electrode coverage into volume space and visualize brain region responses directly on anatomical slices, the FreeSurfer parcellation scheme (aparc + aseg) was utilized. These locations were then grouped together structurally into 28 brain regions to reduce dimensionality and visualized on the MNI152 atlas (**S1 Table, S3 Fig**).

#### sEEG preprocessing

Preprocessing was performed via custom scripts implemented in MATLAB R2021a. DC offset was first removed from raw sEEG signals using a fourth-order Butterworth highpass filter with a cutoff frequency at 0.5 Hz. Then, channels with excessive 60 Hz line noise, defined as having 60 Hz power exceeding three mean absolute deviations above the median, were identified and excluded from future analysis. Data was re-referenced using common average referencing (CAR) before applying a 60 Hz notch filter to remove line noise.

Baseline data was then extracted and broken into five-second epochs to match the duration for vibrotactile stimulation trials, facilitating a baseline subtraction normalization approach. Stimulation data was also broken into epochs specific to each vibration frequency. To minimize effects of interictal epileptic activity, trials with a spike exceeding 500 µV (absolute value) were excluded from future analysis (**S4 and S5 Figs**).

#### Spectral coherence analysis

All coherence analyses were also conducted via custom scripts implemented in MATLAB R2021a. Spectral coherence was computed across four canonical frequency bands – theta (4-8 Hz), alpha (8-13 Hz), beta (13-30 Hz), and broadband gamma (70-170). To expedite processing time during coherence calculations, the data was downsampled to 500 Hz. Coherence was computed using the magnitude-squared coherence function in MATLAB (mscohere) with a window size of 2.5 seconds and 70% overlap.

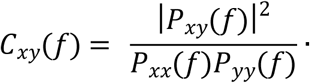

*P_xy_* represents the cross power spectral density (PSD) of *x* and *y*, and *P_xx_* and *P_yy_* represent the PSDs of *x* and *y*, respectively. To normalize the data and facilitate group-level analyses, we performed a baseline subtraction on all coherence values.

#### Signal-to-noise ratio

To identify the vibration frequency/frequencies that elicited the most consistent coherence increase across subjects, we defined a signal-to-noise ratio (SNR), *SNR* = μ/σ. SNR was computed by calculating the mean, *µ*, and standard deviation, *σ*, of the responder percentages for each subject across all five vibration frequencies.

### 2.6 Statistical analysis

Nonparametric statistical analyses were performed in MATLAB R2021a and RStudio. Wilcoxon signed-rank tests were used to assess for statistical significance of group average coherence changes. Based on the hypothesis that vibrotactile taVNS would increase coherence across relevant brain regions, one-sample, right-tailed tests were used to identify distributions significantly greater than 0. The significance threshold was corrected for multiple comparisons (Bonferroni-corrected, 5 vibration conditions tested) and the effect size was computed as *r* = *Z/√N*, where *Z* represents the *Z*-statistic and *N* represents the sample size. Responder pairs across each vibration frequency for individual subjects were identified as those with a Cohen’s d greater than 0.2 when comparing stimulation to baseline.

At the individual subject level, Friedman tests for repeated measures analysis of variance were used to compare MSC during vibration to baseline across all five vibration frequencies. When indicated by a significant result, post-hoc Wilcoxon signed-rank tests (Bonferroni-corrected) were used to identify the significant frequency condition(s). This allowed for identification of which frequencies resulted in the strongest global coherence responses for each subject.

## 3 Results

### 3.1 Global coherence analysis

To assess for brain-wide coherence changes during vibrotactile taVNS, the group-average coherence change from baseline across all possible brain region pairs was aggregated together into a global distribution (n = 353 pairs). Wilcoxon signed-rank tests (Bonferroni-corrected, 5 comparisons) were performed to assess whether the global coherence significantly increased from baseline. Vibrotactile taVNS significantly increased MSC in the theta band as compared to baseline when administered at vibration frequencies of 6, 20, and 40 Hz (6 Hz: *p* = 1.23 x 10^-8^, effect size r = 0.311; 20 Hz: *p* = 7.35 x 10^-9^, effect size r = 0.316; 40 Hz: *p* = 1.82 x 10^-6^; effect size r = 0.264) (**Fig 2**). Alpha coherence was also significantly increased during 6, 20, and 40 Hz vibration (6 Hz: *p* = 0.045, effect size r = 0.126; 20 Hz: *p* = 2.88 x 10^-17^, effect size r = 0.455; 40 Hz: *p* = 1.11 x 10^-13^, effect size r = 0.402). Beta coherence exhibited marginal increases during 6 and 40 Hz (6 Hz: *p* = 0.010, effect size r = 0.153; 40 Hz: *p* = 2.09 x 10^-3^, effect size r = 0.178). No significant changes were observed for broadband gamma coherence (all *p* > 0.05).

**Fig 2.**
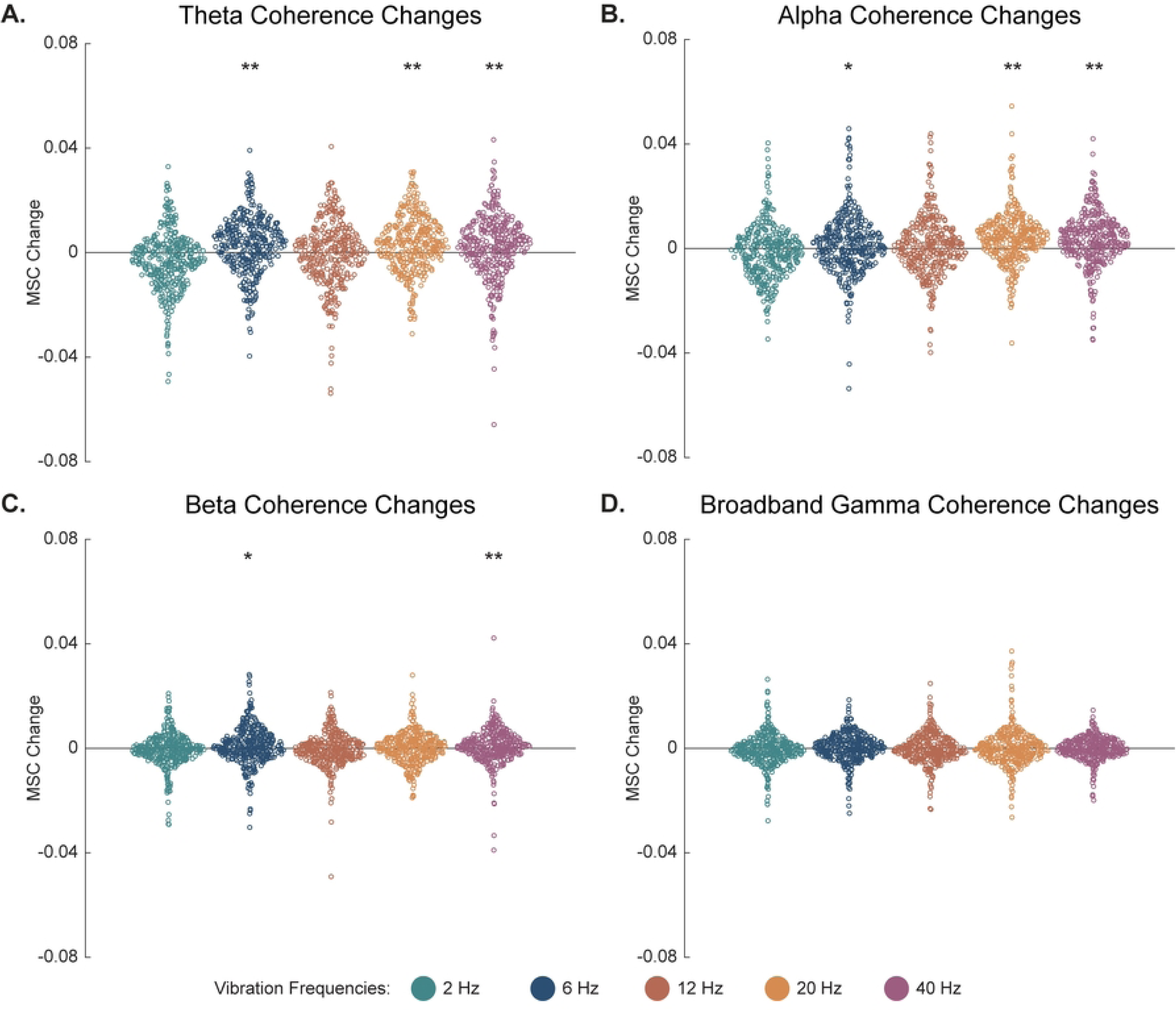
Group-level distributions of magnitude-squared coherence (MSC) changes across theta (A), alpha (B), beta (C), and broadband gamma (D) physiological frequency bands. MSC changes are compared for five vibration frequencies – 2, 6, 12, 20, and 40 Hz. Each point in the MSC distributions represents the coherence change for a particular pair of anatomical regions. Significance threshold Bonferroni-corrected for multiple comparisons. One-tailed, one-sample Wilcoxon signed-rank test: * = Bonferroni-corrected p < 0.05; ** = Bonferroni-corrected p < 0.01.

### 3.2 Subject-specific frequency analysis

To assess which vibration frequency condition(s) subjects were responding to most strongly, we established a responder criterion. Within a subject-specific distribution of all possible electrode pairs, a pair was classified as a responder if the Cohen’s d for stimulation compared to baseline was greater than 0.2. To facilitate comparisons across subjects, we then converted the total number of responder pairs per subject to a percentage of total pairs. Our findings demonstrate that there is subject specificity in terms of optimal vibration frequency (**Fig 3**). Regarding theta coherence, two subjects had the highest responder percentage to 6 Hz vibration, one subject to 12 Hz vibration, three subjects to 20 Hz vibration, and one subject to 40 Hz vibration (**Fig 3A**). For alpha coherence, subject variability was less, with four subjects exhibiting the highest responder percentage to 20 Hz vibration and three subjects to 40 Hz vibration (**Fig 3B**). We also computed SNR (**Section 2.5.4**) for each vibration frequency and then ranked them in descending order to quantify response consistency. The top two frequencies that elicited the most consistent theta coherence increase were 40 and 20 Hz, while 20 and 6 Hz elicited the most consistent alpha coherence increase.

**Fig 3.**
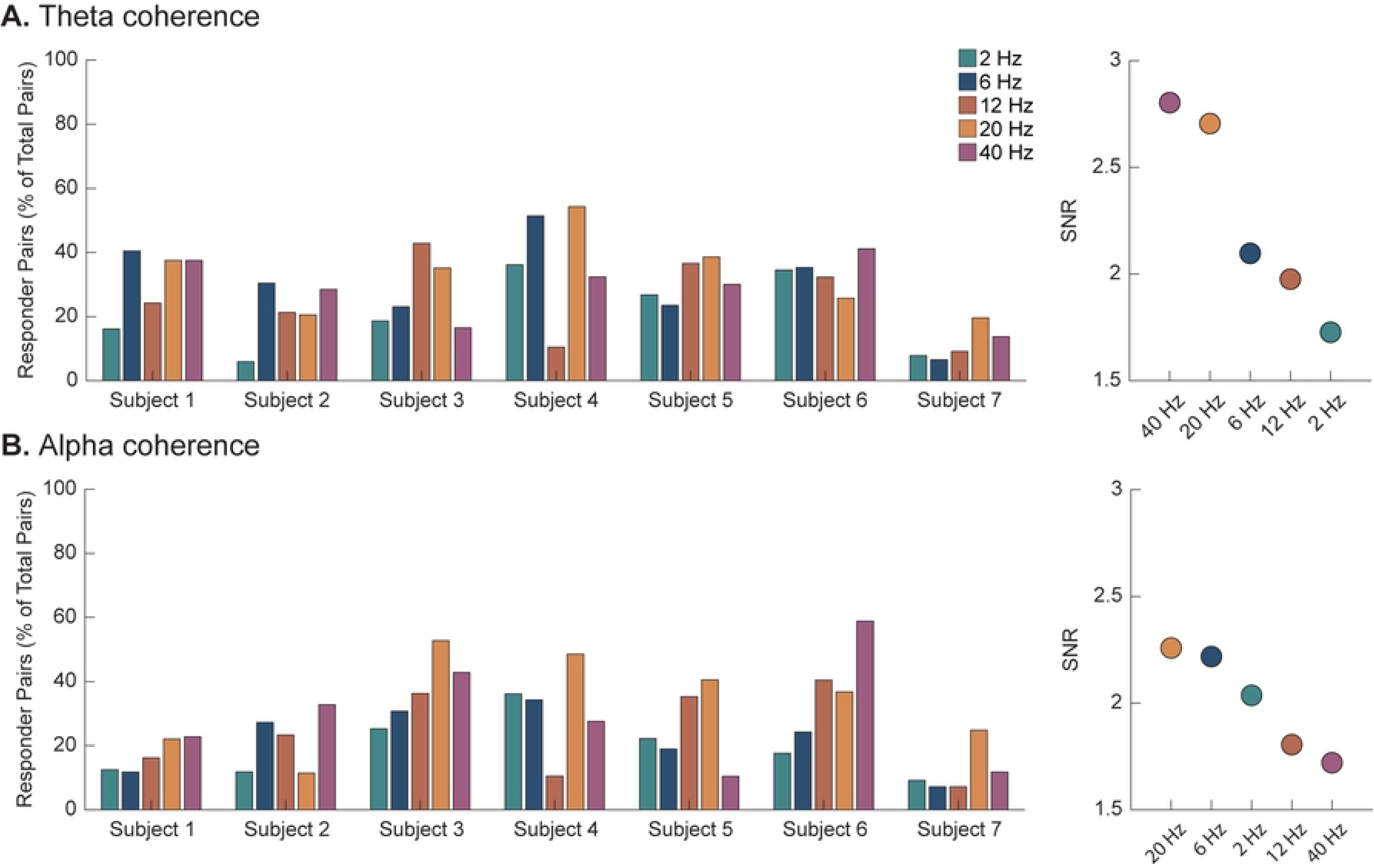
Responder pairs for individual subjects in the theta (A) and alpha (B) frequency bands for all vibration conditions. Responders are defined as pairs of electrodes with a Cohen’s d greater than 0.2 when comparing the mean response during stimulation vs. baseline. Subject specificity is present as subjects have unique vibration conditions that elicit the greatest number of responses. A signal-to-noise ratio (SNR) was computed for each vibration frequency based on the individual subject measurements to generate an overall ranking at the group level.

### 3.3 Exemplar subject

Due to the demonstrated subject specificity (**Fig 3**), we also analyzed individual subjects at the electrode level. An exemplar subject is shown based on bilateral coverage of electrode placement overlapping with all five predetermined ROIs (**Fig 4**). As demonstrated in the coherence maps, this subject had strong coherence increases in both theta and alpha during 40 Hz vibration, as well as a theta increase during 6 Hz vibration (**Fig 4**). A Friedman test was implemented to compare the region-specific coherence values across all five vibration frequencies against the baseline, revealing significant differences between the conditions (theta: χ^2^ = 219.164, *p* = 2.244 x 10^-45^; alpha: χ^2^ = 149.321, *p* = 1.863 x 10^-30^). Post-hoc Wilcoxon signed-rank tests were then conducted to determine which vibration frequencies significantly altered coherence (**S6 Fig**). Theta coherence was significantly increased during 6 and 40 Hz vibration and significantly decreased during 2 Hz vibration (all *p* < 0.001). Specific to alpha coherence, results indicated a significant increase during 40 Hz vibration and a significant decrease during 2 and 20 Hz vibration (all *p* < 0.001).

**Fig 4.**
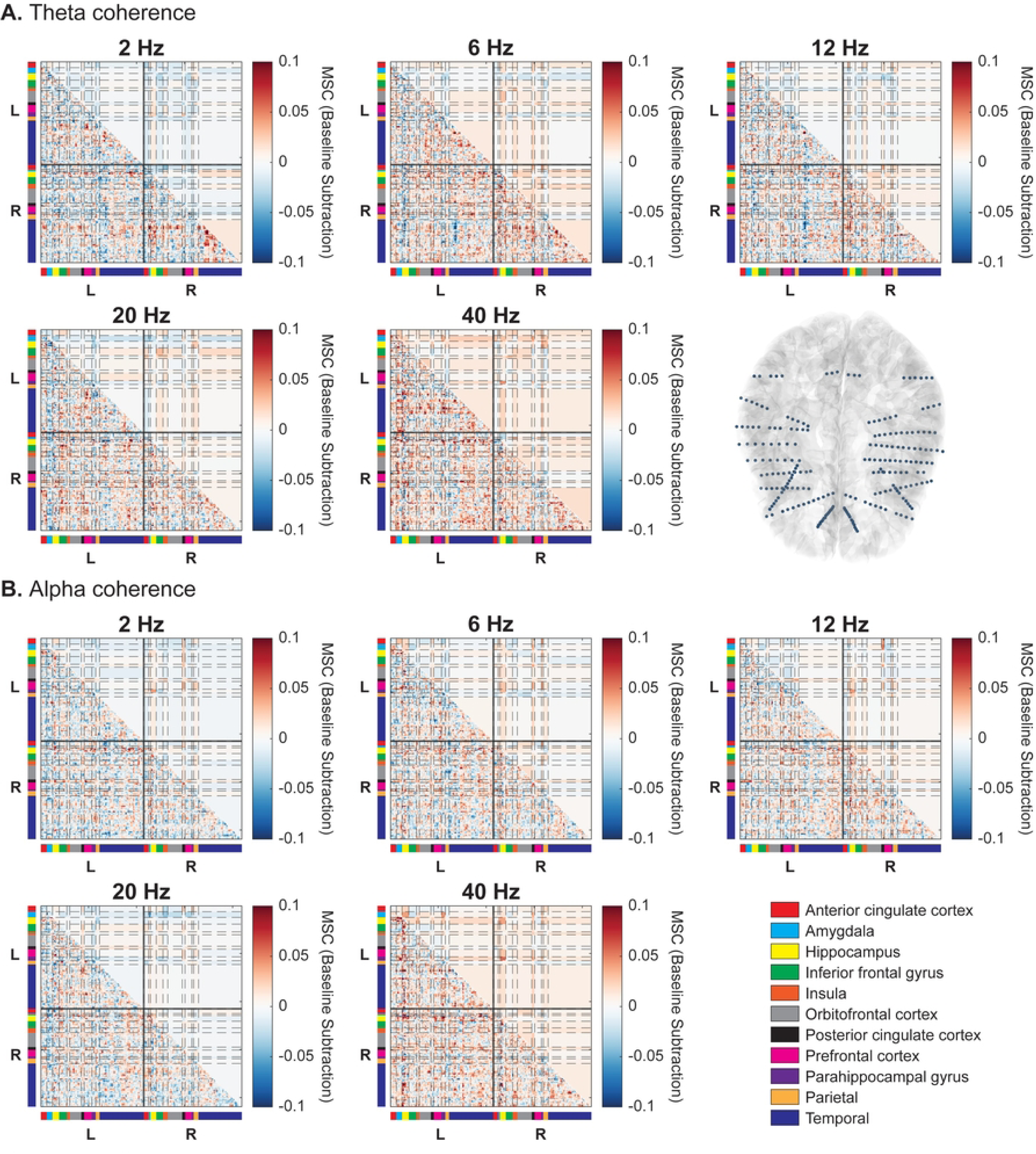
Exemplar subject magnitude-squared coherence (MSC) in the theta (A) and alpha (B) frequency bands for all anatomical regions. The selected exemplar subject had comparable bilateral electrode coverage across both hemispheres (as depicted in the bottom right of **A**). The lower triangle illustrates the coherence matrix for all individual electrodes, whereas the upper triangle shows the region-average coherence matrix.

### 3.4 Data-driven responders

We then took a data-driven approach to identify which electrode pairs exhibited the strongest coherence changes during vibration. As demonstrated in **Fig 2**, vibration frequencies of 6, 20, and 40 Hz resulted in statistically significant global coherence increases in the theta and alpha bands.

Electrode pairs with a coherence increase of ≥ 2 standard deviations above the brain-wide mean were identified (**Fig 5**). These pairings correspond to the points in the top portion of the brain-wide distributions above (**Fig 2**). Brain regions with significant coherence increases to two or more brain regions were left ACC (theta – 6, 20 Hz; alpha – 6, 20 Hz), left PFC (alpha – 6 Hz), left amygdala (alpha – 6 Hz), right ACC (alpha – 20 Hz), right BG (theta – 20 Hz; alpha – 20 Hz), right PHG (theta – 6, 20, 40 Hz; alpha – 6, 40 Hz), right posterior cingulate cortex (PCC) (alpha – 6, 20, 40 Hz), and right central cortex (theta – 40 Hz; alpha – 20 Hz). Complete group-average connectivity matrices can be found in **S7 Fig**.

**Fig 5.**
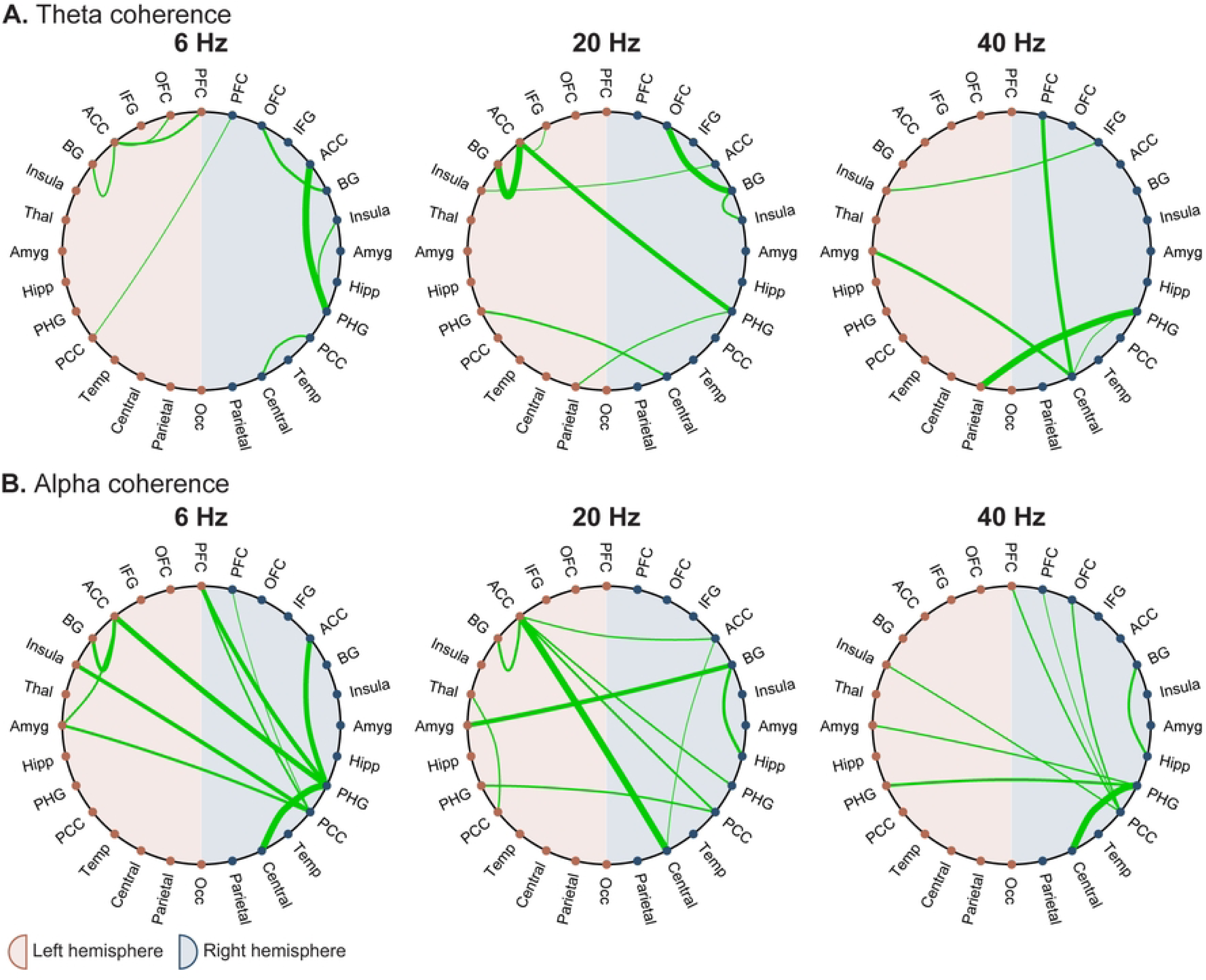
Theta (A) and alpha (B) coherence pairs that are at least 2 standard deviations above the mean for 6, 20, and 40 Hz vibration conditions as identified in Fig 2. Only coherence pairs that are ≥ 2 standard deviations above the mean are shown, and the line thickness is relative to the magnitude, such that the thickest line in each plot is the pair with the greatest coherence increase. Note that the line thickness is relative to the particular plot (i.e., each pairing with the thickest line does not have the same magnitude coherence increase; rather, each pairing with the thickest line represents the greatest coherence increase shown in that plot). All coherence pairings can be found in **S7 Fig**. Left regions are denoted with red points while right regions are denoted with blue points. Brain regions are sorted from anterior to posterior when looking from top to bottom of the left or right hemisphere. ACC = anterior cingulate cortex; Amyg = amygdala; BG = basal ganglia; Hipp = hippocampus; IFG = inferior frontal gyrus; Occ = occipital lobe; OFC = orbitofrontal cortex; PCC = posterior cingulate cortex; PFC = prefrontal cortex; PHG = parahippocampal gyrus; Temp = temporal lobe; Thal = thalamus.

### 3.5 Region- and frequency-specific responses

Predetermined ROIs were defined as seeds to test the hypothesis that vibrotactile taVNS would result in increased coherence along the vagal activation pathway. Theta and alpha coherence exhibited anatomically distinct changes during vibrotactile stimulation compared to baseline (**Fig 6**).

**Fig 6.**
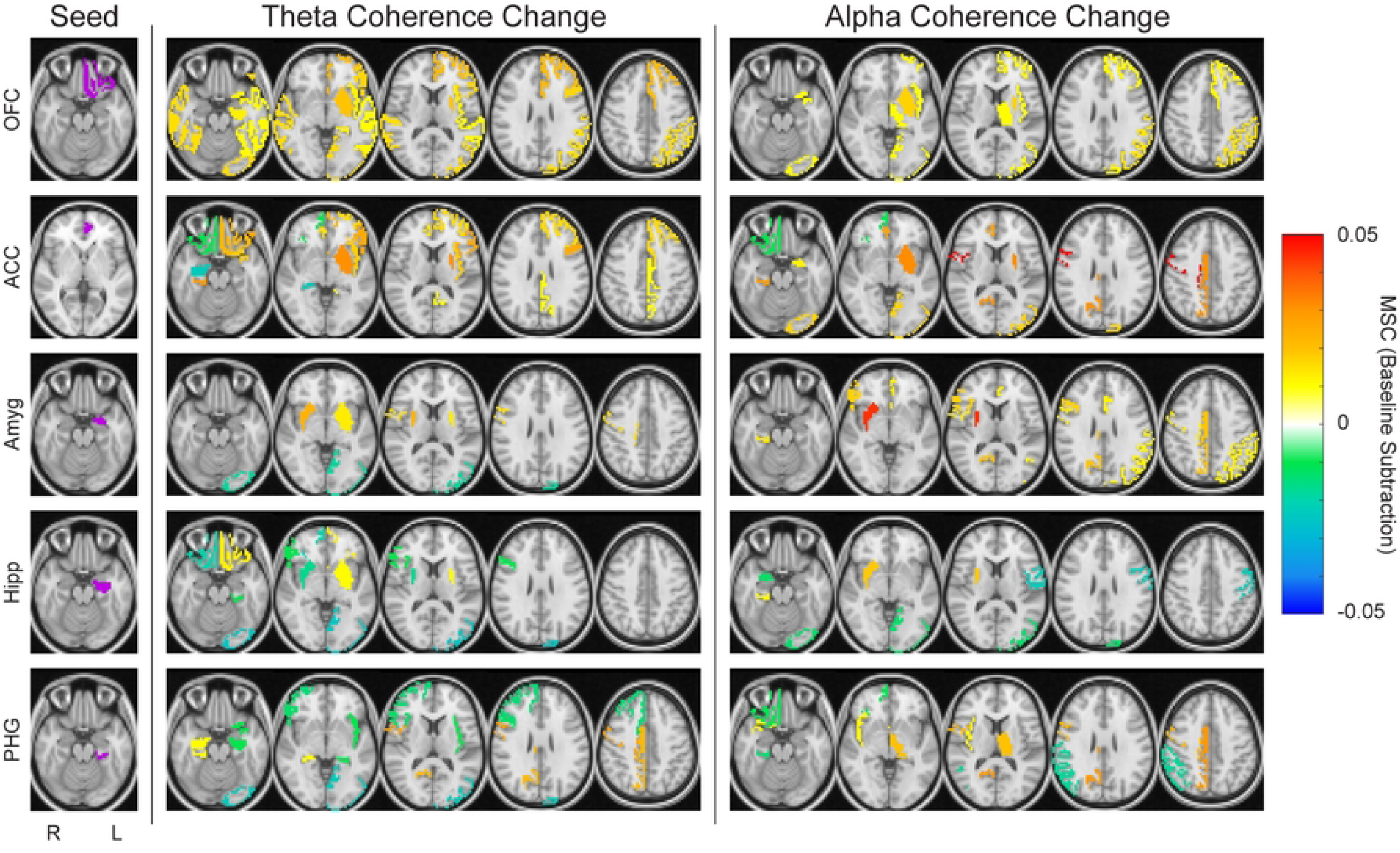
Coherence changes to identified regions of interest (ROIs) at the group level. ROIs were set as the seed region, and coherence changes are shown to the rest of the regions encompassed in the sEEG coverage. A threshold of 0.01 was set such that only changes with an absolute value > 0.01 are shown. Both theta and alpha coherence changes are shown, all under the 20 Hz vibration condition. Increased coherence during vibration is shown as a positive coherence change (warm tones), while decreased coherence during vibration is shown as a negative coherence change (cool tones). From top to bottom, seeds are left orbitofrontal cortex (OFC), left anterior cingulate cortex (ACC), left amygdala (Amyg), left hippocampus (Hipp), and left parahippocampal gyrus (PHG). Images are shown in radiological view. The highlighted seed regions are mapped based on predetermined groups of FreeSurfer parcellations to reduce dimensionality (see **S1 Table and S3 Fig**).

Specifically, each anatomic seed (left OFC, left ACC, left amygdala, left hippocampus, and left PHG) had distinct anatomic coherence changes. Coherence changes varied depending not only on the anatomic seed, but also on the physiological frequency band. While theta and alpha coherence exhibited some anatomic overlap for a given seed, there were also notable differences. As an example, with 20 Hz stimulation both the left OFC and ACC seeds demonstrated a notable increase in theta and alpha coherence with the ipsilateral BG. However, the left amygdala seed demonstrated a broad alpha coherence increase with the ipsilateral parietal lobe that was not present in the theta band.

Conversely, to isolate effects specific to the vibration frequency parameter, the anatomic seed was held constant while comparing coherence patterns. This analysis similarly resulted in diverse anatomic patterns of theta and alpha coherence dependent on the vibration frequency condition (**Fig 7**). Also notable is that the different vibration conditions elicited different anatomic distributions. As an example, the anatomic distribution of theta coherence for the left amygdala seed was very different between 2 and 40 Hz stimulation, with regions such as the right BG and ACC exhibiting responses in opposite directions.

**Fig 7.**
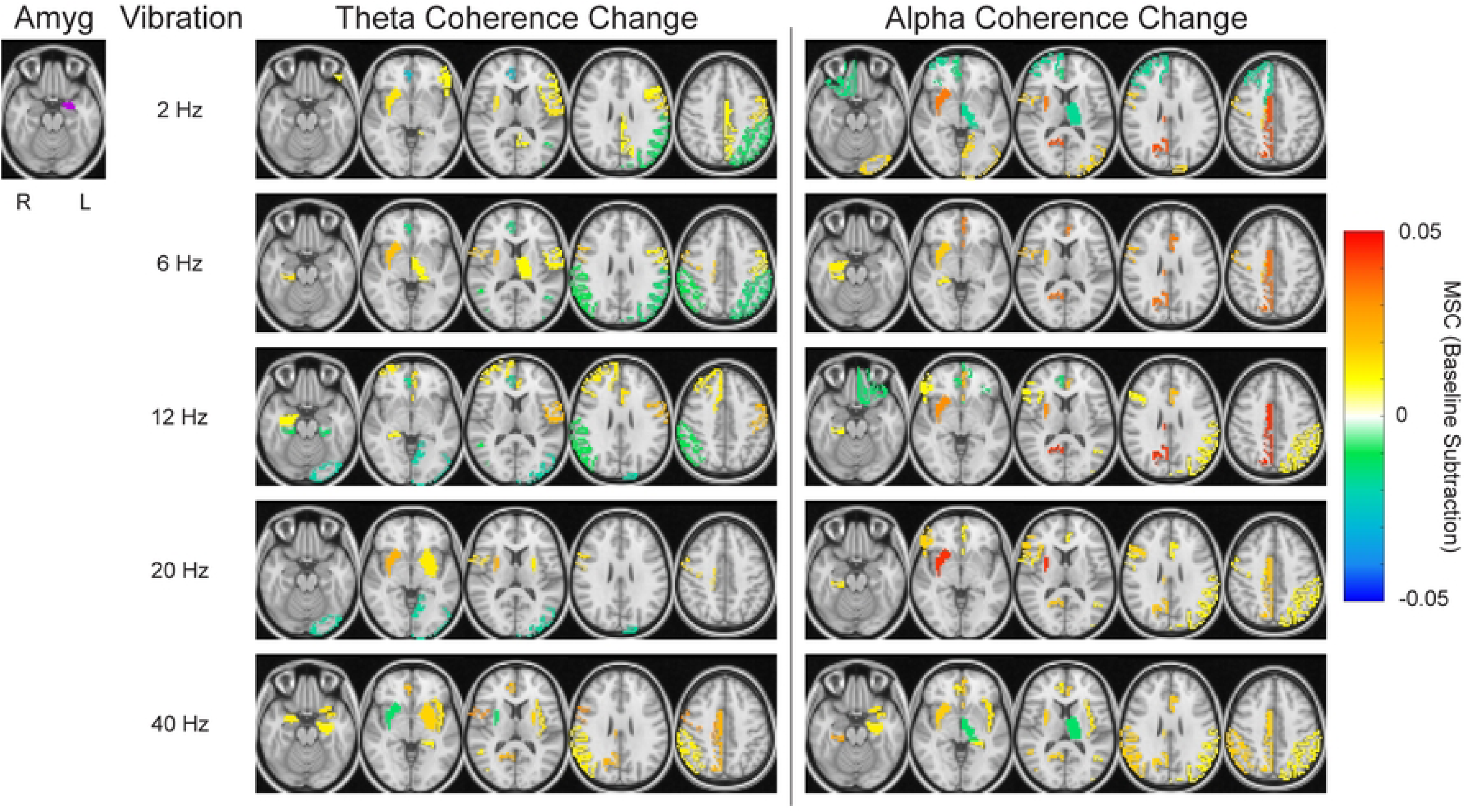
Theta and alpha coherence changes across all five vibration conditions for a specific seed region of interest. The left amygdala was chosen as the seed due to all subjects having electrode coverage in that region. As in Fig 6, a threshold of 0.01 was set, and increased coherence during vibration is shown as a positive coherence change, while decreased coherence during vibration is shown as a negative coherence change. Images are shown in radiological view.

## 4 Discussion

We investigated the potential of vibrotactile stimulation administered to the cymba concha region of the outer ear as an alternative modality for taVNS by investigating resulting neuronal coherence. To do so, we characterized neurological responses to vibration at varying frequencies in subjects undergoing continuous sEEG monitoring. This invasive human physiology served to validate and provide insights into the noninvasive methodology. Specifically, the present work demonstrates that vibrotactile taVNS activates key brain regions purported to underly mechanism(s) of action for VNS and taVNS.

### 4.1 Prior work of vibrotactile taVNS

To our knowledge, there have been two prior studies that investigated the use of vibration as an alternative taVNS modality. The first study sought to extend known anti-inflammatory effects of invasive VNS to a vibrotactile modality [62,63]. Addorisio and colleagues quantified the effect of vibrotactile ear stimulation on inflammatory markers in healthy adults and assessed its utility from a clinical efficacy standpoint in patients with RA, revealing beneficial anti-inflammatory effects [46]. While direct comparisons cannot be drawn between the present study and this work, it provided the first evidence supporting the use of vibration as an alternative to electrical stimulation to activate the ABVN. Another study demonstrated the potential of 6 Hz vibration administered to the cymba concha to improve working memory performance, while also highlighting the ability of vibrotactile taVNS to increase general arousal during the course of repeated working memory tasks [47].

In addition to the aforementioned studies specific to vibrotactile stimulation of the cymba concha in humans, prior anatomical studies also help lay the foundation for our hypothesis that vibrotactile taVNS can activate vagal pathways. Specifically, the afferent makeup of the ABVN lends support to the hypothesis that vibration could activate those fibers. ABVN fibers are somatic afferents, indicating they are sensitive to touch and pressure and thus theoretically vibration as well [64,65]. Of note, the cymba concha, the stimulation site in this study, is innervated solely by the ABVN [66]. What has been lacking to date, however, is a more thorough characterization of the central impact of vibrotactile stimulation to the ABVN, which this study provides.

### 4.2 Group-level characterization of vibrotactile taVNS in humans

At the group level, our results suggest an overall increase in global low-frequency (theta and alpha) coherence, most evident during 6, 20, and 40 Hz vibration (**Fig 2**). Studies have shown that tasks requiring enhanced arousal and higher cognitive load, such as spatial conflict or working memory tasks, are associated with increases in theta and alpha coherence [67–70]. Of note, central vagus nerve projections, such as those activated by VNS and taVNS, influence arousal [71–75]. While the mechanisms for our observed parametric specificity in terms of stimulation frequency are not clearly understood, we hypothesize that the rhythm structure of the sensory input may have different resonances with underlying neural circuitry. The specificity of responses depending on vibration frequency complements current literature on electrical VNS and taVNS studies, which have demonstrated frequency-specific effects [55,76–79]. Additionally, the anatomic locations of cortical and subcortical regions that demonstrated the most robust coherence changes are structurally distinct. Thus, it is consistent that frequency rhythms known to be associated with longer-range interactions (i.e., theta, alpha) would demonstrate preferential effects [80,81].

To anatomically resolve the regions that demonstrated enhanced coherence, we performed a data-driven analysis to classify sites with the most robust responses to stimulation (**Fig 5**). Overlapping with our initial hypothesis, the ACC and PHG were key regions that emerged as hubs. These findings align with rodent models by Cao et al. (2016) that revealed an increased theta band correlation between the basolateral amygdala and ACC following VNS [39].

We further analyzed brain-wide responses within theta and alpha coherence in predefined cortical and subcortical ROIs. Several limbic ROIs exhibited increased coherence to each other, and the thalamus and PCC emerged as other sites with increased coherence to multiple ROIs (**Fig 6**). The limbic system and thalamus have been well-studied in the context of VNS [38,82,83]. Our results demonstrate that they have enhanced connectivity, which supports the notion that vibrotactile taVNS activates central structures similar to classic electrical VNS/taVNS. Of note, the coherence effects across these ROIs varied depending on whether theta or alpha rhythms were assessed. ROI coherence was also dependent on vibration frequency (**Fig 7**). Parametric analyses of VNS have shown the importance of parameter selection in terms of both physiological and behavioral responses, and our work is consistent in illustrating that vibration frequency selection is also relevant in the context of vibrotactile taVNS. A multitude of factors may explain this parametric sensitivity of neuronal responses. The diverse projections stemming from the vagus nerve and the variable proximity of other nuclei to the vagus nerve termination site may lead to distinct temporal sensitivities [55].

### 4.3 Subject-specific frequency responses to vibrotactile taVNS

Individualized responses to neuromodulation techniques and broader instances of personalized medicine are well-documented in the literature [84–89]. Accordingly, we also assessed variance in response to the different vibrotactile parameters across individuals. For the purposes of this study, we defined the optimal frequency as that which resulted in the greatest percentage of electrode pairs classified as responders. While the same three frequencies emerged through this analysis as at the group level (6, 20, and 40 Hz), inter-subject variability was present in terms of which of those frequencies resulted in the most widespread coherence increases (**Fig 3**). However, it is important to note that inter-subject response variability may be somewhat attributable to the inter-subject variability in electrode coverage (**Fig 1**). As an example, we highlighted the MSC changes in Subject 2, who had near-symmetrical bilateral coverage (**Fig 4**). Despite the individual response variation across vibration frequencies, there were vibration parameters that elicited more consistent central responses. Specifically, 20 Hz vibration frequency had a high SNR for inducing both theta and alpha coherence increases. For theta specifically, however, 40 Hz stimulation had a slightly higher SNR. Moving forward, we propose using 20 Hz vibrotactile taVNS to increase global low-frequency coherence as it maximizes the SNR across theta and alpha and had the highest effect size for increased global coherence in those bands. Alternatively, to customize vibrotactile taVNS and achieve maximum coherence increases in an individual patient, further studies are needed to identify and validate noninvasive means for selecting a patient’s optimal stimulation parameters.

### 4.4 Limitations

To collect valuable invasive brain recordings from human subjects, data was recorded from patients with intractable epilepsy undergoing invasive monitoring via sEEG. These recordings offer critical insights into how cortical and subcortical structures such as the amygdala and hippocampus respond to vibrotactile taVNS. However, we note that data from a clinical cohort may not be indicative of wider populations. Future studies with other clinical populations or healthy controls using noninvasive neuroimaging modalities (e.g., fMRI, EEG) may enhance the reproducibility and translatability of our present findings. Further, the study is limited by the sample size; however, given the novelty of vibrotactile taVNS, the goal was to determine whether there is a central response to justify continued research. An additional limitation of this work concerns the sound produced by vibration motors. While administering vibrotactile stimulation to the outer ear, the stimulation caused both mechanical and auditory perception. Future work should investigate whether auditory stimulation alone without mechanosensation results in similar coherence changes in the brain.

## 5 Conclusions

Vibrotactile taVNS offers an alternative approach to activating vagal fibers to influence central brain dynamics. The present work leveraged human intracranial recordings to demonstrate the viability of vibrotactile taVNS to enhance coherence between central cortical and subcortical structures known to be associated with classic electrical VNS stimulation. Further, the central effects are subject-specific and can be variable depending on both the frequency band assessed and the frequency of vibrotactile stimulation.

## Acknowledgements

The authors would like to thank the clinical team on the Epilepsy Monitoring Unit at Barnes Jewish Hospital for supporting data collection efforts. We also thank all the patients who participated in this study.

## 9 Supporting information captions

**S1 Table. Aggregated FreeSurfer groups.** FreeSurfer parcellations were grouped structurally into 28 defined brain regions to reduce the dimensionality. ACC = anterior cingulate cortex; Amyg = amygdala; BG = basal ganglia; Hipp = hippocampus; IFG = inferior frontal gyrus; OFC = orbitofrontal cortex; Occ = occipital lobe; PCC = posterior cingulate cortex; PFC = prefrontal cortex; PHG = parahippocampal gyrus; Temp = temporal lobe; Thal = thalamus

**S1 Fig. Study design schematic.** Each experimental session consisted of 3 minutes of baseline followed by a 25-minute stimulation period. During that time, stimulation alternated between 5 seconds on and 5 seconds off, with randomized vibration frequencies being delivered. In total, each vibration frequency was delivered for 30 trials, totaling 150 trials across all frequencies.

**S2 Fig. Custom earpiece used for vibrotactile taVNS.** The 3D-printed earpiece is shown from a front view **(A)**, back view **(B)**, and angled view **(C)**. The stimulating end is indicated by a solid line while the stabilizing end is indicated by a dashed line. The stimulating end contacts the cymba concha, and the vibration motor is inserted into the slot visible in the back view. The stabilizing end wraps around the back of the ear for support. **(D).** Image of the device positioned on the ear.

**S3 Fig. All FreeSurfer parcellations included in anatomical analysis.** ACC = anterior cingulate cortex; Amyg = amygdala; BG = basal ganglia; Hipp = hippocampus; IFG = inferior frontal gyrus; Occ = occipital lobe; OFC = orbitofrontal cortex; PCC = posterior cingulate cortex; PFC = prefrontal cortex; PHG = parahippocampal gyrus; Temp = temporal lobe; Thal = thalamus

**S4 Fig. Rejecting trials with interictal epileptiform discharges. (A)** All preprocessed baseline epochs (5 s) from one electrode in one subject. Any trial with a positive or negative spike exceeding abs(500) µV (two trials highlighted here in teal and pink) was marked as containing epileptiform discharges and excluded from future analysis. **(B)** This example electrode is located in the right amygdala.

**S5 Fig. Proportion of total trials rejected per subject according to methodology detailed in S4 Fig.** For each subject, all electrodes are considered in aggregate, for a total trial number of 30 *x* total electrode count for baseline, and 30 *x* total electrode count *x* 5 vibration conditions for vibration. The percentage of trials included for analysis is shown in the lighter color, while the percentage of rejected trials is shown in the darker color for both baseline (dark blue) and vibration conditions (teal).

**S6 Fig. Exemplar subject magnitude-squared coherence (MSC) in the theta (A) and alpha (B) frequency bands.** Distributions are directly compared between baseline and stimulation conditions. Red asterisks indicate that the global coherence distribution is significantly decreased during stimulation for this exemplar subject, while green asterisks indicate that it is significantly increased during stimulation. Wilcoxon signed-rank test: ** = p < 0.001.

**S7 Fig. Group-average magnitude-squared coherence (MSC) in the (A) theta and (B) alpha frequency bands for all anatomical regions.** MSC plots are shown for 6, 20, and 40 Hz vibration conditions as identified in **Fig 2**. Because anatomical coverage is clinically driven, there is variation in the number of subjects represented within each coherence voxel. Voxels with an MSC change equal to 0 are pairings that do not exist across any subject. ACC = anterior cingulate cortex; Amyg = amygdala; BG = basal ganglia; Hipp = hippocampus; IFG = inferior frontal gyrus; Occ = occipital lobe; OFC = orbitofrontal cortex; PCC = posterior cingulate cortex; PFC = prefrontal cortex; PHG = parahippocampal gyrus; Temp = temporal lobe; Thal = thalamus.

